# ImmunoFusion: A Unified Platform for Investigating RNA-seq-Derived Gene Fusions in Cancer and Immunotherapy

**DOI:** 10.1101/2025.06.05.658082

**Authors:** YanKun Zhao, Shixiang Wang, Shensuo Li, Minjun Chen, Su-han Jin, Udo S. Gaipl, Hu Ma, Jian-Guo Zhou

## Abstract

**Background:** Gene fusions play a critical role in cancer development by persistently activating kinases or inactivating tumor suppressor genes, leading to altered signal transduction and gene expression regulation. However, their impact on treatment responses remains poorly understood. Although existing cancer databases catalog numerous fusion events or immune checkpoint blockade (ICB) studies, no unified platform integrates gene fusion data across cancer types while linking them to the tumor microenvironment (TME) and patient outcomes. Such integration is essential for elucidating how fusions shape immune responses and for developing improved biomarkers for personalized cancer therapies.

**Methods:** To address this gap, we constructed the Fusion Immune Atlas (ImmunoFusion), a platform integrating data from TCGA, TARGET, CPTAC, and 29 ICB cohorts (including ICB treatments and other treatment modalities) across diverse cancer types. We identified fusion events from raw RNA-seq data using Arriba and STAR-Fusion, and standardized fusion calls by adapting MetaFusion to the GRCh38 reference genome. Additionally, we curated clinical data and estimated TME signatures and cell fractions using the Immuno-Oncology Biological Research (IOBR) approach. ImmunoFusion was developed using the high-quality Rhino framework for Shiny applications, built on R.

**Results:** ImmunoFusion (https://shiny.zhoulab.ac.cn/ImmunoFusion) encompasses 21,014 clinical samples, serving as a comprehensive RNA-seq-derived gene fusion database and analytical tool. It offers functionalities to investigate fusion breakpoints, confidence scores, frequency patterns, and associations with clinical variables. The platform also enables TME evaluation and interactive exploration of fusions for analyzing tumor immunophenotypes in cancer and immunotherapy contexts. As an illustration, analysis of *MTAP* fusions in lung cancer cohorts revealed their association with a metabolically depleted, “cold” (immune-suppressive) tumor environment and poorer patient outcomes, pinpointing *MTAP* fusions as a novel biomarker for treatment selection.

**Conclusions:** ImmunoFusion represents a significant advancement, delivering a unified platform to explore the influence of gene fusions on cancer and immune responses. It offers particular value in understanding fusion-driven immunotherapy outcomes, paving the way for more effective therapeutic strategies.

## Introduction

Oncogenic gene fusions have transformed cancer diagnostics and therapeutics. Over the past century, from the pivotal role of *BCR*::*ABL1* in chronic myeloid leukemia to *ALK*, *ROS1*, and *RET* fusions guiding targeted therapies in non-small cell lung cancer (NSCLC), these chimeric products drive carcinogenesis by aberrant transcription or generating oncoproteins^1^. An analysis of available data shows that gene fusions occur in all malignancies, and that they account for 20%^2^, but their functional heterogeneity is driven by complex genomic instability, necessitates transcriptomic validation to differentiate driver events^3, 4^ from biological noise (e.g., physiological chimeric RNAs^5^, readthrough transcripts^6^, conjoined/duplicated genes^7, 8^ and tandem arrangements^4^). Recent evidence suggests that distinct fusion events may cooperatively define molecular subtypes, as seen with *RET* and *NTRK* fusions in pediatric thyroid cancers^9^. Existing gene fusion databases primarily catalog key fusion genes or events rather than comprehensively scanning fusions, thus failing to fully address the complexity of cancer gene fusions.

RNA sequencing (RNA-seq) offers superior sensitivity and resolution over traditional methods for detecting gene fusions, yet it poses challenges in prioritizing pathogenic fusions among millions of candidates. Accurate identification of fusion isoforms in the transcriptome and distinguishing driver fusions are critical for precision oncology. Arriba^10^ and STAR-Fusion^11^, two established fusion detection tools, have been widely applied to large cancer cohorts, including The Cancer Genome Atlas (TCGA). Large-scale fusion databases, including TumorFusions^12^ and ChimerDB4.0^13^, primarily integrate TCGA RNA-seq data to identify fusion candidates across cancer types, but most rely on a single fusion-calling algorithm. Given frequent discrepancies among callers, integrated tools combining their strengths are essential for improved accuracy^14^. Moreover, well-known projects, such as the Clinical Proteomic Tumor Analysis Consortium (CPTAC) and Therapeutically Applicable Research to Generate Effective Treatments (TARGET), lack dedicated fusion databases, restricting their utility for fusion analysis.

Immune checkpoint blockade (ICB) has transformed cancer treatment, yet many tumors remain unresponsive, with durable benefits limited to a small subset of patients. Current biomarkers, such as PD-L1^15^ and tumor mutation burden (TMB)^16^, provide limited predictive accuracy, underscoring the need for novel biomarkers derived from other molecular alterations. Although gene fusions are established oncogenic drivers, their role in predicting ICB response remains largely underexplored. Recent studies have found that *ALK* fusions are associated with poor ICB response in NSCLC patients^17^, whereas *CDK12* fusions correlate with improved ICB response in prostate cancer^18^. However, no systematic platform currently exists to investigate the interplay between gene fusions and ICB response.

To address these unmet needs, we developed ImmunoFusion, a comprehensive database and analysis platform integrating RNA-seq-derived gene fusions from TCGA, TARGET, CPTAC, and 29 ICB cohorts. By integrating fusion calls from Arriba and STAR-Fusion using MetaFusion, ImmunoFusion creates, to our knowledge, the largest cancer gene fusion database. This platform facilitates exploration of interactions among fusion events, clinical features, and the TME, enabling systematic identification of gene fusions as biomarkers for cancer prognosis and treatment response. By elucidating the role of gene fusions in cancer and the TME, ImmunoFusion enhances our understanding of molecular alterations and advances insights into immuno-oncology.

## Results

### Construction of the cancer gene fusion catalogue

ImmunoFusion (accessible at https://shiny.zhoulab.ac.cn/ImmunoFusion/) was developed through a streamlined data processing pipeline encompassing data collection, preprocessing, and integration. We collected TCGA, TARGET, and CPTAC data (N=16,324) from the GDC data portal and 29 ICB cohorts with RNA-seq data sourced from genome sequence archive repositories (N=4,690) (Figure 1A, B; see Methods). Gene fusions were detected from normal (N=1,807) and tumor samples (N=19,267) using two top-performing algorithms Arriba^10^ and STAR-fusion^11^ to detect RNA-seq fusion calls^19^. We utilized MetaFusion^20^ to standardize the outputs of Arriba and STAR-Fusion, identifying 69,836 fusions in normal samples and 586,607 fusions in tumor samples. Following rigorous preprocessing (see Supplementary Methods), we identified 2,097 high-confidence (confident) fusions in normal tissues and 93,310 confident fusions in tumor tissues (Figure 1B). Clinical data, patient outcomes, and cancer and TME signatures were cleaned and standardized to enhance cross-cohort consistency and comparability. ImmunoFusion encompasses 39 cancer types and provides a unified database and analysis platform to facilitate exploration of gene fusions in cancer and ICB (Figure 1C, D). Unlike existing databases^12, 13, 21–31^, which primarily focus on fusion events, annotations, or ICB cohorts (Supplementary Table 1), ImmunoFusion integrates these elements comprehensively.

**Figure 1.**
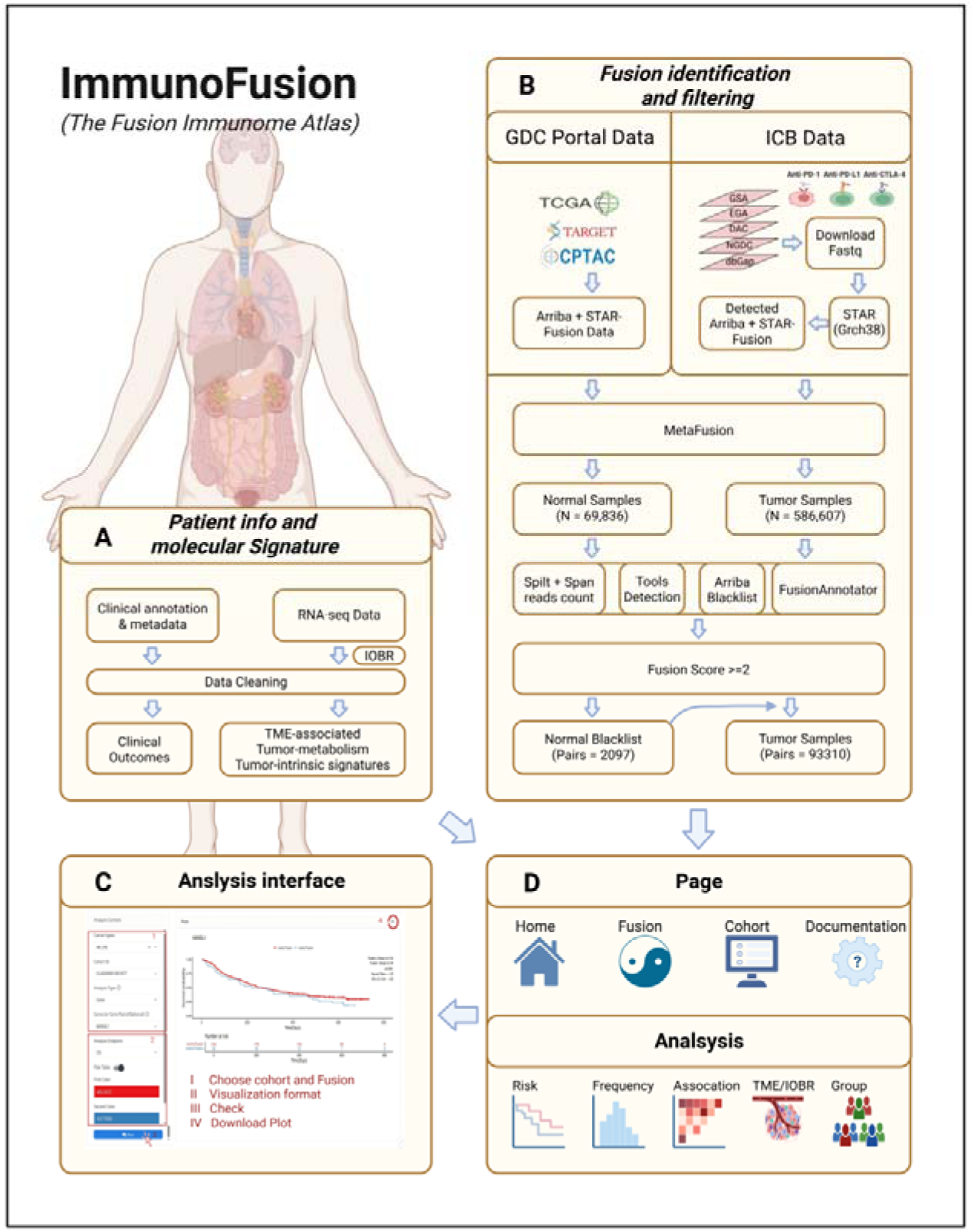
Overview of ImmunoFusion. (**A**) Workflow for acquiring and processing patient data, cancer signatures, and tumor microenvironment (TME) signatures. (**B**) Pipeline for identifying, integrating, and scoring gene fusions. (**C**) Example analysis page on the ImmunoFusion platform, illustrating analysis steps and results, with options to save results as PDF or PNG images in user-specified dimensions. (**D**) ImmunoFusion offers four web pages and five analytical modules, including survival analysis, cancer and TME signatures, fusion frequency, fusion associations, and group analysis.

### Statistics of ImmunoFusion

The distribution of gene fusions is categorized as follows (Figure 2A): (1) Sample type: normal samples yielded 114,483 fusions, while tumor samples yielded 986,816 fusions. (2) Fusion type: MetaFusion classified fusions into six subtypes (Supplementary Figure 1): CodingFusion (normal: 54.7%, tumor: 53.9%), ReadThrough (15.8%, 18.9%), TruncatedNoncoding (14.1%, 13.4%), TruncatedCoding (12.8%, 11.6%), SameGene (2.2%, 2.1%), and NoHeadGene (0.3%, 0.2%). (3) Fusion confidence score: A stringent scoring system was implemented to evaluate reliability of fusion candidates (Supplementary Methods), with approximately 81.9% of fusions scoring below 2, defined unconfident in this study and thus filter them out for further exploration. We annotated unique fusions using FusionAnnotator, revealing that 96.5% of fusions in normal samples and 18.4% in tumor samples lacked annotations (Figure 2C, D). We observed high variability in fusion abundance, measured as record counts, across cancer types (Figure 2B).

**Figure 2.**
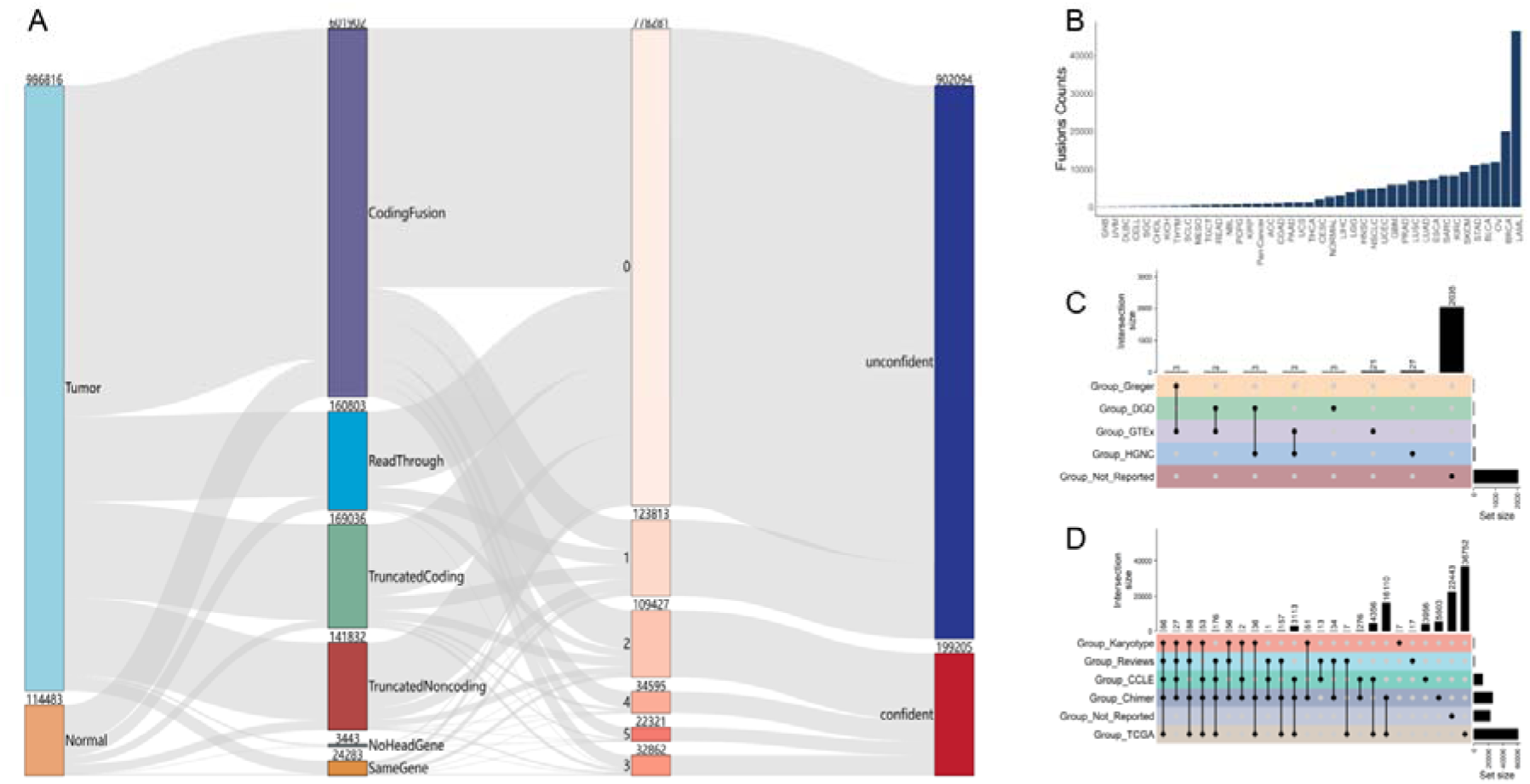
Distribution of MetaFusion integrated gene fusions. **(A)** Sankey diagram illustrating the stratification of fusion records counts. The leftmost node represents sample types. The second-layer node displays fusion classifications by MetaFusion based on 5’/3’ gene breakpoints and coding region annotations. Subsequent layers depict filtration steps in the ImmunoFusion workflow, with the final layer categorizing fusion pairs as confident (score ≥2, high-confidence) or unconfident (score <2, low-confidence). **(B)** Number of confident cancer fusions across cancer types. Note: Pan-cancer cohort detail() **(C)** Distribution of confident fusions in normal samples annotated by FusionAnnotator, potentially unrelated to cancer or representing false positives. Horizontal bars (right) indicate fusion counts per annotation. Dots and lines represent fusion subsets. Vertical histograms show gene counts per subset. Group_Greger: tandem genes from Greger *et al.*^32^; Group_HGNC: HGNC gene family membership; Group_GTEx: recurrent fusions in GTEx normal samples detected by STAR-Fusion^11^; Group_DGD: Duplicated genes from Ouedraogo *et al.*^8^; Group_Not_Reported: confident fusions not reported previously identified in normal samples. **(D)** Distribution of confident fusions in tumor samples annotated by FusionAnnotator. Horizontal bars (right) denote fusion counts per annotation. Group_Karyotype: Known fusions from the Mitelman Database^27^; Group_Reviews: 2019 COSMIC reported fusions^33^ + literature-curated cancer fusions by FusionAnnotator; Group_CCLE: Fusions in DepMap cancer cell lines^34^ + Klijn *et al.* study^35^ + TCGA cell line data (STAR-Fusion)^11^; Group_Chimer: ChimerDB-reported fusions^36^; Group_Not_Reported: Unannotated confident cancer fusions; Group_TCGA: Fusions from DEEPEST-Fusion^37^ + Guo *et al.*^14^ + TumorFusions ^12^ + TCGA RNA-seq (STAR-Fusion)^11^ + Alaei-Mahabadi *et al.*^38^ + YOSHIHARA *et al.*^3^.

To provide a comprehensive overview of gene fusion frequency and the contribution of dominant fusions across cancer and sample types, we constructed a pan-cancer landscape of confident gene fusions and their original cohort resources (Figure 3A-C). Our data reveal distinct patterns of fusion burden across cancer types, with significant variability in sample size and fusion burden. Hematologic malignancies like LAML (N=2,772; 46,601 fusions) exhibit near-universal fusion positivity (98.7%) but low per-sample fusion rates (∼17/sample), suggesting reliance on a limited set of driver events. In contrast, solid tumors such as stomach adenocarcinoma (STAD) (N=457; 11,072 fusions; ∼25/sample) and ovarian serous cystadenocarcinoma (OV) (N=428; 11,844 fusions; ∼23/sample) demonstrate disproportionately high fusion burdens despite smaller cohort sizes—likely driven by genomic instability (OV’s propensity for structural variations^39^, STAD’s complication in histological and etiological heterogeneity^40^). Consistent with prior reports^14, 41^, kidney renal clear cell carcinoma (KIRC) (N=2,006; 8,317 fusions; ∼5/sample) exhibits a markedly lower fusion burden.

**Figure 3.**
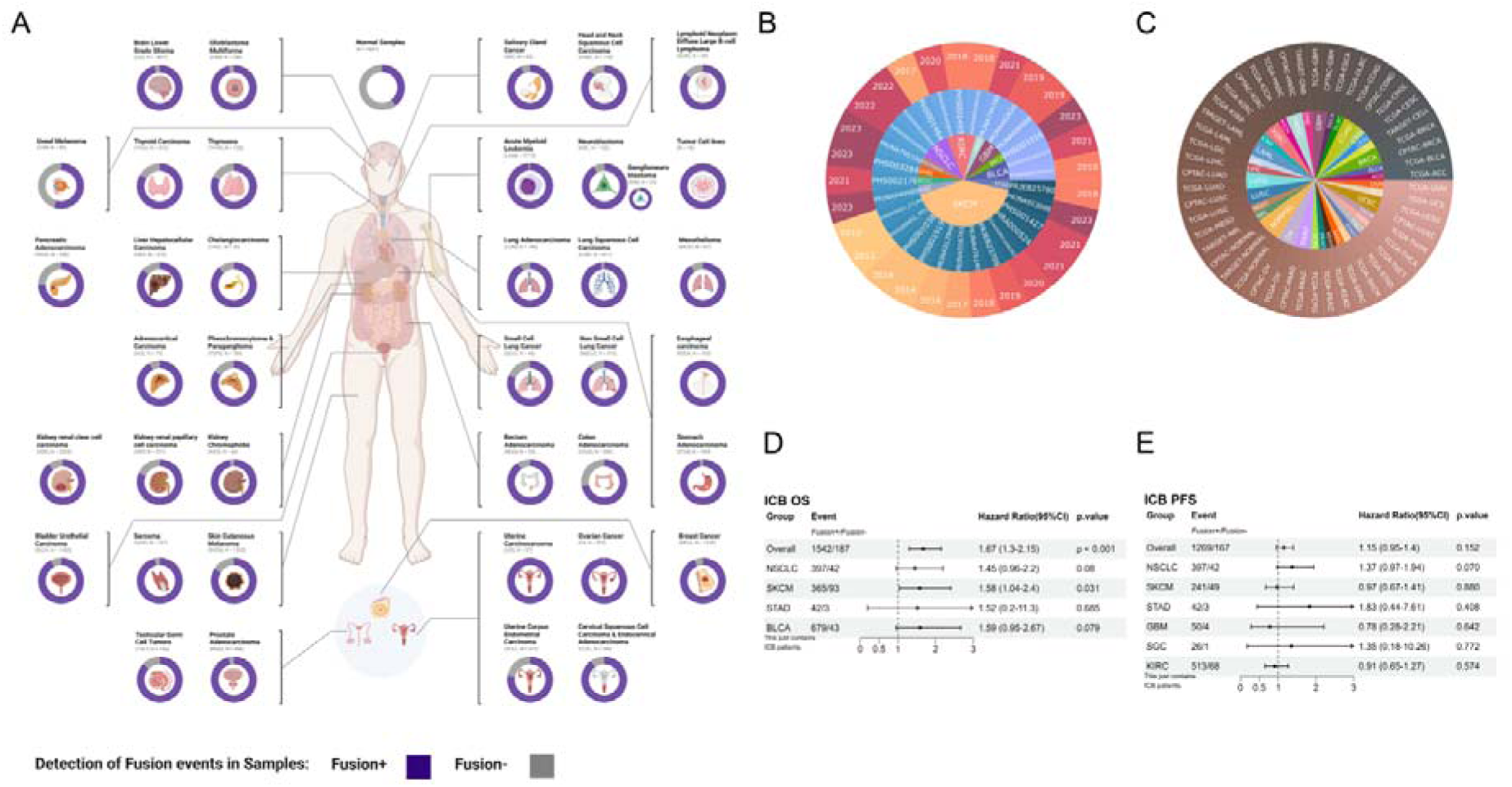
Data landscape of ImmunoFusion. **(A)** An overview of confident gene fusion events detected in each cancer type or normal samples. For a given cancer type, subtypes are indicated with a ring around the corresponding inset organ view. Breaks in each ring distinguish whether fusions were detected in the samples. (The Pancer-cohort^42^ was excluded from visualization due to unavailability of tumor type annotations for samples and its predominant reliance on blood transcriptomes data; Normal samples from patients with diverse cancer types were consolidated into a single category and represented as a circle; tumor cell lines, derived from the TARGET-AML project cell line subset, were visualized separately.) **(B, C)** Inclusion of cancer types, cohort and publication years in this study between ICB cohort to GDC. **(D, E)** These forest plots illustrate the hazard ratios (HR) and 95% confidence intervals (CI) for tumor-type-stratified analyses of OS and PFS in the Cox proportional hazards model, restricted to ICB-treated patients with pre-treatment samples.The aggregate of all cohorts is denoted as “Overall”. Associations with p < 0.001 are indicated as ‘p < 0.001’.

### Gene Fusions associated survival landscape

For ICB treated subgroups, in IPD meta-analysis, while fusion-detected (Fusion+) patients exhibited significantly shorter overall survival (OS) (HR=1.67_Fusion+ vs Fusion-_ [1.3-2.1], p<0.001; Median=19.1 months(m) vs 27 m) (Figure 3D), no statistical difference was observed in progression-free survival (PFS) (p=0.152) (Figure 3E). In tumor type-specific analyses, Fusion+ cases in NSCLC showed a trend toward reduced overall survival (OS; HR=1.45 [95% CI: 0.96–2.20], p=0.08) and progression-free survival (PFS; HR=1.37 [95% CI: 0.97–1.94], p=0.07). In SKCM, Fusion+ cases were associated with significantly worse OS (HR=1.58 [95% CI: 1.04–2.40], p=0.031), but PFS showed minimal change (p=0.880). Similar trends were observed across other tumor types.

Similarly, GDC IPD meta-analysis revealed significantly elevated risks for both OS (HR=1.48 [1.28-1.7], p<0.001) (Supplementary Figure 2A) and PFS (HR=1.23 [1.08-1.4], p=0.002) (Supplementary Figure 2B) in Fusion+ patients. GDC data also revealed cancer specific divergent prognostic impacts: colon adenocarcinoma (COAD) Fusion+ patients showed worse OS and PFS. In comparison, skin cutaneous melanoma (SKCM) Fusion+ patients exhibited improved OS and PFS; Lung adenocarcinoma (LUAD) displayed non-significant OS differences (Supplement Figure 2A) but favorable PFS (HR=0.63 [0.42-0.95], p=0.027) (Supplement Figure 2B); LAML(HR=0.51 [0.27-0.95], p=0.033; Median=17.9 vs 8.9) (Supplement Figure 2A); neuroblastoma(NBL) (HR_Fusion+ vs Fusion-_=0.34 [0.14-0.86], p=0.022; Median=54.5 vs 10.8) (Supplement Figure 2A) demonstrated significant OS benefits in Fusion+ patients **(Other cohort details in** Supplement Figure 2A, B). These findings underscore context dependent and cancer specific biological roles of fusions, necessitating integrated functional annotation and TME profiling to elucidate underlying mechanisms.

### Web functionality of ImmunoFusion

ImmunoFusion offers exploratory and advanced analytical tools, enabling researchers to explore complex relationships and gain actionable insights (Figure 1D). Modules are organized across five pages: Home, Fusion, Cohort, Analysis, and Help. Users can explore associations between fusion events, clinical variables, TME features, molecular signatures, therapeutic responses, and prognosis. Researchers can identify fusions tied to prognosis or immunotherapy response and validate findings across cohorts or tumor types. A standardized, user-friendly workflow is provided (Figure 1C). Below, we detail the core modules.

*Fusion* page: The ImmunoFusion *Fusion* page offers a comprehensive resource for exploring and retrieving gene fusions identified across cancer cohorts. It integrates essential details such as gene symbols, genomic positions (hg38), cohort of origin, fusion type, breakpoint exon regions, confidence metrics (including ImmunoFusion score, prior reporting status, and read counts), and distribution patterns across cohorts. Users can easily locate specific fusions using search filters based on gene symbols or any text for regular expression matching.

*Cohort* page: The *Cohort* page provides an intuitive platform for accessing datasets, allowing users to filter cohorts by origin, cancer type, treatment modality, therapeutic agents, and cohort size. Detailed cohort-level metadata is presented in comprehensive tables, facilitating informed cohort selection.

*Analysis* page: The *Analysis* page enables multi-tiered exploration of fusion events at the gene, fusion, and record levels, facilitating in-depth investigation of their roles in tumor progression, immune modulation, and immunotherapy outcomes. This framework identifies differential biological impacts of distinct fusion partners, revealing potential biomarkers. Key sub-modules include: *Distribution*, which employs stacked bar plots to display fusion gene frequencies across tumor types and cohorts, highlighting specific or pan-cancer prevalence; *Comparison*, which identifies significant changes in TME cell type infiltrations or 255 cancer-associated signatures—categorized into TME-related, tumor metabolism, and tumor-intrinsic groups—associated with specific fusion events; *Association*, which uses Fisher’s exact test to generate heatmap plots visualizing co-occurrence or mutual exclusivity of fusion events, elucidating their correlations; *Kaplan-Meier (KM) and Cox Analysis*, where KM generates survival curves for comparative studies and Cox evaluates prognostic associations of fusions; and *Group Analysis*, which enables nuanced comparisons of fusion events across multiple groups within a cohort, uncovering intricate patterns through dual-mode analysis.

### Case studies validating and extending data of MTAP Fusion

Emerging evidence establishes *MTAP* expression which represents the most frequently homozygously deleted region^43^ as an independent prognostic factor with greater clinical significance than *CDKN2A* in NSCLC^44^. Recent multi-cancer analyses demonstrate that combined *CDKN2A*/*MTAP* loss correlates with immunologically “cold” TME characterized and inferior responses to ICI patient^45^. However, these studies did not investigate *MTAP* fusion events.

MTAP fusions primarily result from deletion events, with detailed fusion characteristics from Arriba provided in Supplementary Table 2. As a use case, we utilized ImmunoFusion to investigate MTAP fusion events in NSCLC cohorts.

In the POPLAR cohort (N = 192; EGAD00001008548), *MTAP* fusion (2.6% incidence, Figure 4A) predicted reduced survival (OS HR _MTAP+ vs MTAP-_=2.52 [0.63-10.07, p=0.034; 6.4 vs 11.4; Figure 4B). Similar patterns emerged in the OAK cohort (N=699; EGAD00001008549) with 1.6% fusion frequency (Figure 4D) showing worse outcomes (OS HR=2.05 [0.85-5.05], p=0.02; Median=6.4 vs 11.5; Figure 4E; PFS HR=1.86 [0.83-4.18], p=0.035; Median=2.2 vs 2.9; Figure 4F). TCGA-LUAD cohort (N=506; 3.1% incidence, Figure 4G) confirmed the poor OS outcome (HR=1.01 [0.83-5.66] p=0.020; Median=26.9 vs 50; Figure 4H), though PFS lacked significance (Figure 4I). Furthermore, to explore differential treatment effects in NSCLC, we performed treatment-stratified analyses, In the atezolizumab-treated subgroups (Supplementary Figures 3A, B, E, F), the POPLAR cohort demonstrated significantly elevated OS risk (HR=3.2, p=0.015, Median=5.7 vs 14.5; Supplementary Figures 3A]), although not statistically significant, the OAK cohort showed a similar tendency (OS HR=2.3; Median=9.4 vs 12.8). Meanwhile, the OAK docetaxel-treated subgroup exhibited significantly inferior OS (HR=1.87; Median=4.7 vs 10.7; Supplementary Figure 3I). *MTAP* fusion patients tend to have nonresponse (Supplementary Figure 3C, D, G, H). *MTAP* Fusion+ NSCLC is associated with progression following atezolizumab monotherapy versus docetaxel monotherapy and tend to result in primary resistance to anti-PD-L1 therapy, as indicated by the lack of response in the majority of Fusion+ cases.

**Figure 4.**
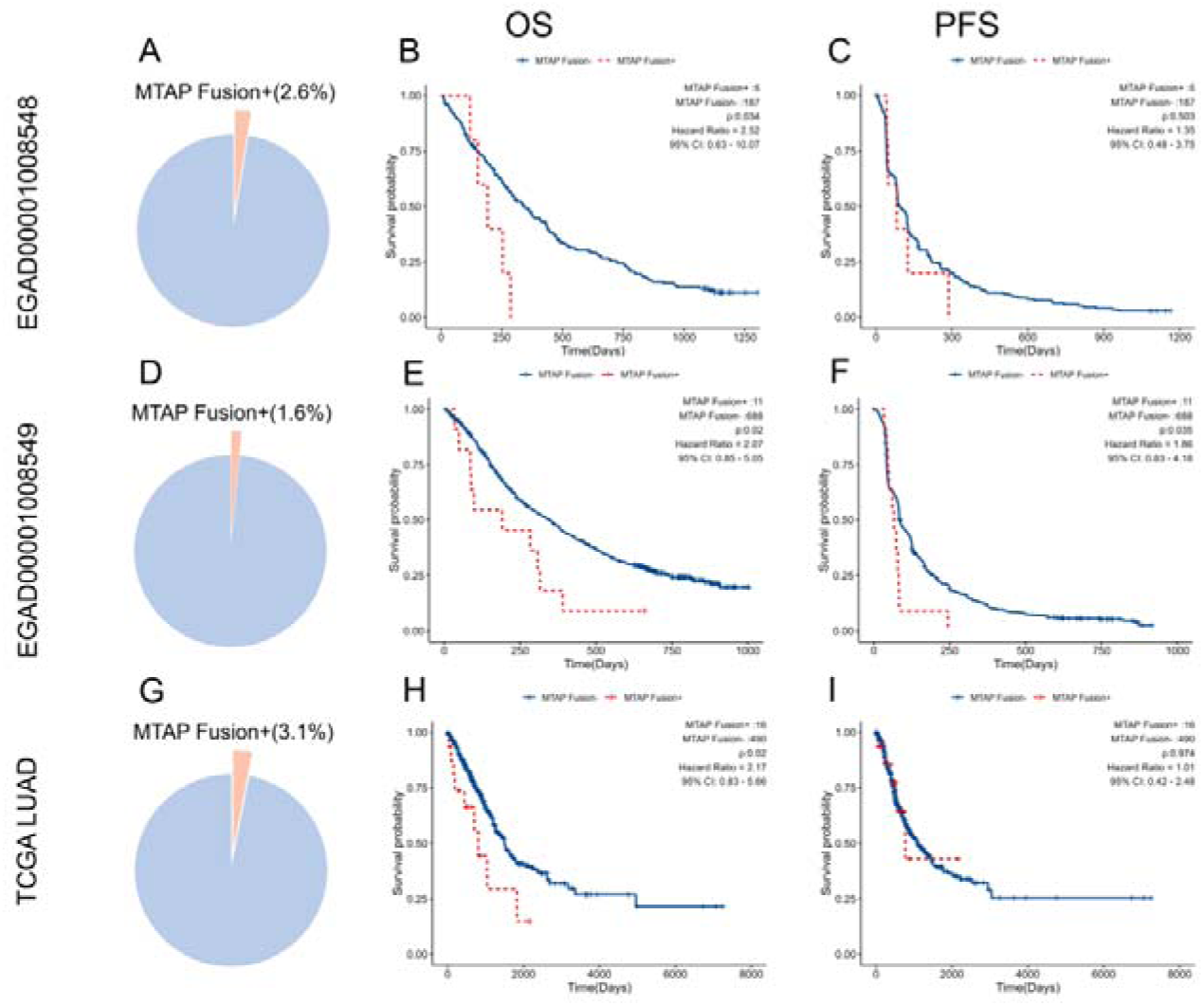
MTAP fusion prevalence and prognostic impact in NSCLC cohorts (Pail et al(POPLAR, OAK)^51^, TCGA-LUAD). **(A, D, G)** Pie charts depict the proportion of MTAP fusion-detected (Fusion+: orange) and fusion-not detected (Fusion−: blue) patients in the POPLAR (N = 192; Fusion+ = 5, 2.6%), OAK (N = 699; Fusion+ = 11, 1.6%), and TCGA-LUAD (N = 506; Fusion+ = 16, 3.1%) cohorts. (B, C, E, F) (Fusion+: red; Fusion-: blue) Kaplan-Meier curves for overall survival (OS) and progression-free survival (PFS) stratified by MTAP fusion status in patients treated with chemotherapy (docetaxel) or anti-PD-L1 therapy (atezolizumab) (POPLAR: B, C; OAK: E, F). (H, I) Survival analyses in the TCGA-LUAD cohort (non-treatment-specific). Hazard ratios (HR) with 95% confidence intervals were calculated using Log-Rank Test.

Mechanistically, loss of MTAP leads to extracellular accumulation of its substrate methylthioadenosine (MTA), a metabolite with potent immunomodulatory effects. Tumor-derived MTA suppresses T-cell function by engaging the adenosine A2B receptor (A2BR)^46–48^, contributing to an immunosuppressive TME. Additionally, MTAP deletion influences the occurrence and progression of tumors through multiple metabolic pathways^49^. Emerging evidence further implicates MTA in broader T-cell dysregulation^50^, fostering a “cold” TME and primary resistance to ICB^45^.

Our multi-cohort TME analysis of *MTAP* fusion-positive (Fusion+) tumors reveals conserved metabolic vulnerabilities with context-dependent immunosuppressive features. Fusion+ tumors exhibit elevated hypoxia signatures (Figure 5A), reflecting a metabolic adaptation to oxygen scarcity (Figure 5A). Enhanced cardiolipin metabolism (Figure 5B), methionine cycle hyperactivity (Figure 5D) and elevated polyamine levels (Figure 5F) further indicate mitochondrial stress. Paradoxically, despite higher tumor antigen release (Figure 5E), Fusion+ tumors display an immune-cold TME characterized by MDSC enrichment (Figure 5G) and reduced MHC Class II expression (Figure 5C), suggesting defective antigen presentation and myeloid-driven suppression override antigen availability. This dichotomy implies that *MTAP* fusions establish a dual barrier to immunity: metabolic suppression of effector T cells combined with defective antigen recognition.

**Figure 5.**
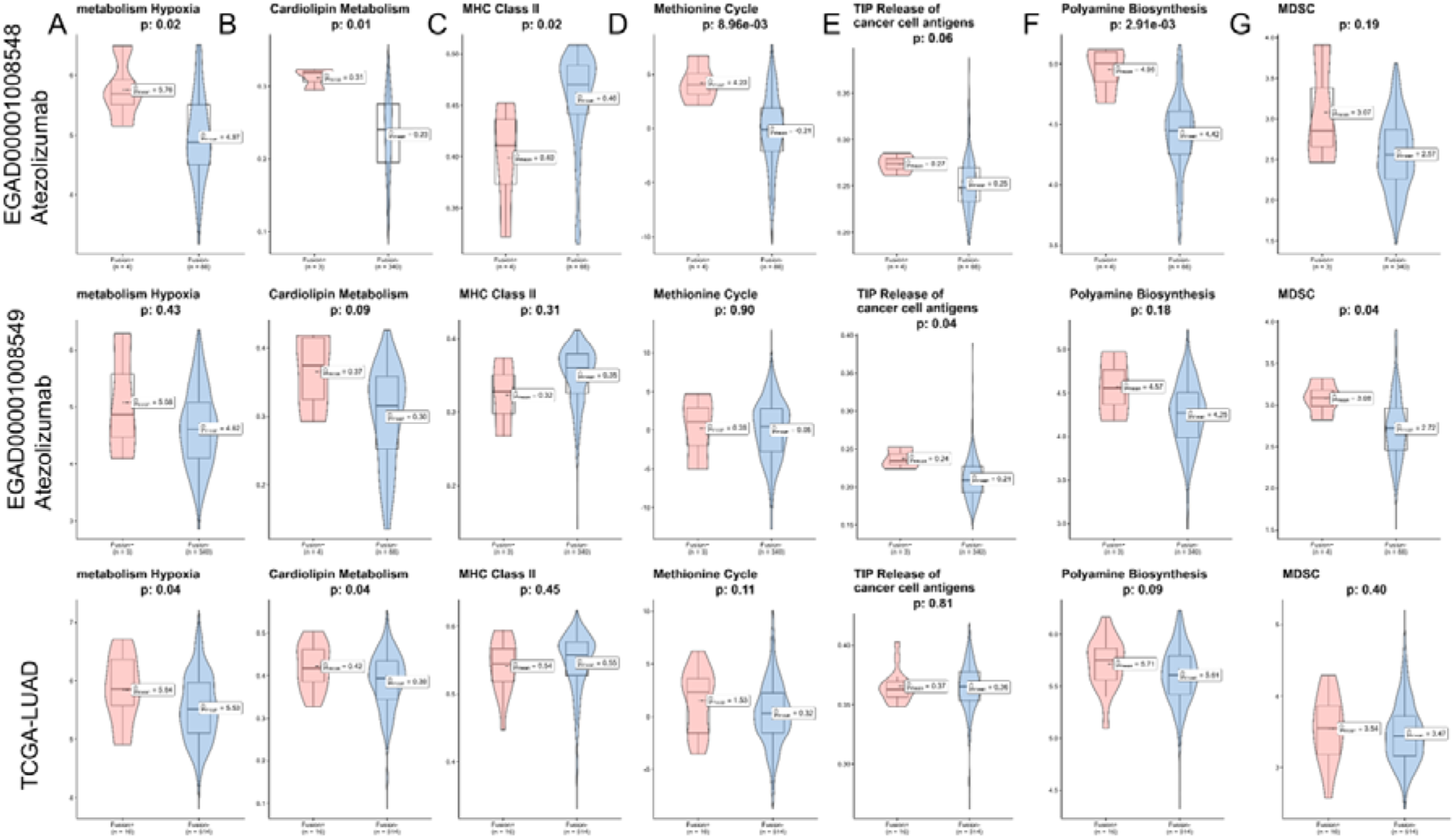
MTAP fusion+ tumors exhibit characteristics of metabolic vulnerability and an immunologically cold TME. **(A) Metabolism Hypoxia (zscore):** Hypoxia scores are elevated in MTAP Fusion+ tumors (EGAD00001008548-Atezolizumab and TCGA-LUAD cohorts). **(B) Cardiolipin Metabolism (ssGSEA):** Mitochondrial dysfunction is enhanced in MTAP Fusion+ tumors (EGAD00001008548-Atezolizumab and TCGA-LUAD cohorts); **(D) Methionine Cycle (PCA):** Hyperactivated methionine metabolism in MTAP Fusion+ tumors (EGAD00001008548-Atezolizumab cohort); **(F) Polyamine Biosynthesis (zscore):** Elevated polyamine levels in MTAP Fusion+ tumors (EGAD00001008548-Atezolizumab cohort); **(C) MHC Class II (ssGSEA):** Paradoxically elevated MHC-II expression in MTAP Fusion+ tumors (EGAD00001008548-Atezolizumab cohort); **(E) TIP Release of cancer cell antigens (ssGSEA):** Increased cancer cell antigen release in MTAP Fusion+ tumors (EGAD00001008549-Atezolizumab cohort); **(G) MDSC (zscore):** MDSC enrichment in MTAP Fusion+ tumors (EGAD00001008549-Atezolizumab cohort); p values were calculated by two-sided Wilcoxon signed-rank test. Number of samples: POPLAR [Fusion+ N=4, Fusion- N=88], OAK [Fusion+ N=3, Fusion- N=341], TCGA-LUAD [Fusion+ N=16, Fusion- N=514]. Boxes indicate the median ± interquartile range; whiskers, 1.5× the interquartile range; centre line, median. Only contains NSCLC patients treated with anti-PD-L1.

## Discussion

Gene fusions are a defining feature in certain cancers^37, 52^, offering substantial potential for clinical research and precision oncology. However, existing frameworks for studying fusion-related cancer biology face critical limitations, including restricted sample sizes and diversity, which hinder the generalizability of findings. Challenges in exploring rare fusions—stemming from functional variability in breakpoint locations, interpretative complexities in fusion detection datasets, and barriers to clinical trial recruitment—further complicate therapeutic prioritization when multiple actionable genetic alterations are present. To address these gaps, we developed ImmunoFusion, an integrated data platform that aggregates fusion events from three large pan-cancer projects and 29 immunotherapy cohorts, encompassing 19,267 cancer samples across 39 cancer types and pan-cancer cohort. ImmunoFusion provides user-friendly visualization, customizable options, and unrestricted access without mandatory registration, making it a cost-effective resource for researchers and clinicians.

In this study, we describe the data sources, collection, and standardization procedures for ImmunoFusion, followed by an overview of its statistics, functionalities and website analysis modules. We also provide a detailed, step-by-step guide to operating the platform. Compared with other tools, ImmunoFusion offers several key advantages. First, it constitutes the largest database for gene fusion exploration in cancer, incorporating previously unavailable data from TARGET and CPTAC. Second, it is the first web server to integrate fusion genes with ICB cohorts, offering enhanced tumor microenvironment characterization compared to databases like FPIA^23^, thereby advancing the study of gene fusions in immuno-oncology. As an example, we systematically analyzed *MTAP* gene fusion events in 1,427 NSCLC patients, revealing their clinical relevance and associations with TME characteristics—findings supported by prior research^45, 53^ on similar mechanisms. By elucidating the interplay between fusion events and the TME and establishing a biomarker exploration framework, ImmunoFusion is poised to become an indispensable resource in ICB research.

Nevertheless, ImmunoFusion faces several data limitations. Sample sizes for certain cancer types, such as BRCA and HNSC, remain limited, and access restrictions to raw RNA-seq datasets^54, 55^ further constrain data comprehensiveness. Integrating data is challenging due to the inherent heterogeneity in fusion events and phenotypic data across cohorts, complicating unified analyses; however, the platform’s flexibility allows users to combine cohorts for customized exploration. Moreover, despite rigorous data processing, integration, and scoring, the detected fusions are computational predictions and lack large-scale laboratory validation. Additionally, ImmunoFusion’s functional scope is focused on ICB, and its analytical cohorts do not include those primarily designed for studying targeted therapies.

We are dedicated to continually enhancing ImmunoFusion by incorporating data from diverse and emerging cohorts in cancer, immunotherapy, and targeted therapies, while integrating new features driven by user feedback. Its distinctive capabilities, robust analytical power, and scalability establish ImmunoFusion as an essential tool for elucidating fusion-driven insights in cancer and informing personalized therapeutic approaches.

## Methods

### Data acquisition from GDC portal

We obtained Arriba and STAR-Fusion results from the Genomic Data Commons (GDC) data portal (https://portal.gdc.cancer.gov/) (accessed on 2023-08-13). Specifically, we downloaded gene fusion datasets from TCGA, TARGET and CPTAC projects. These datasets encompass a total of 14,521 tumor samples and 1807 normal tissues across 36 different tumor types. It is important to note that these cohorts are not specifically related to immunotherapy; rather, they represent publicly available cancer datasets. Their inclusion in our study serves to complement and contrast with the checkpoint immunotherapy cohorts, thereby enabling a more in-depth exploration of the role of fusion genes in tumors.

### Systematic search bulk RNA-seq data from ICB related cohorts

Referring to our previous cohort collection process in TCCIA^56^, we performed a systematic search using PubMed to identify articles related to bulk RNA-seq data from solid cancer patients treated with ICB. In total, we collected 29 studies^51, 57–74^ related to checkpoint immunotherapy that provided raw RNA-seq datasets. Details are described in Supplementary Methods. This analysis included over 4,746 clinical samples from 29 cohorts treated with ICBs including PD-1/PD-L1 and CTLA-4 inhibitors, as well as other treatments. The analysis considered both pre-treatment and on-treatment responses.

### Fusion identification and integration

Raw RNA-seq data were aligned to the human reference genome (GRCh38) using STAR aligner (v2.7.10a). We employed a dual-caller framework, utilizing Arriba (IO: v2.0.0; GDC: v1.1.0, https://github.com/suhrig/arriba) and STAR-Fusion (GDC: v2.0.0; IO: v1.6.0, https://github.com/STAR-Fusion/STAR-Fusion), to identify RNA-derived gene fusion candidates, with results integrated via MetaFusion. A scoring system was developed to assess the credibility of fusions detected in tumor and normal samples. See Supplementary Methods for more details.

### Clinical data preprocessing

We retrieved clinical data from the Genomic Data Commons (GDC) and UCSCXenaShiny ^75^platform, followed by data cleaning and standardization, including filling missing values, removing duplicates, renaming columns, and processing the gender field. Survival, purity, genomic instability, and subtype data were merged using Sample ID. We confirmed that most valid rows in GDC clinical data aligned with those from UCSCXenaShiny, and used UCSCXenaShiny data to impute missing values in the GDC dataset, merging by Sample ID. See Supplementary Methods for more details.

### ImmunoFusion implementation

The ImmunoFusion database is developed as a Web application leveraging R Shiny (https://shiny.posit.co/) and utilized the high-performance and state-of-the-art Rhino (https://appsilon.github.io/rhino/) frame to optimize our Shiny framework through The Appsilon Way. ImmunoFusion, is developed solely for research purposes and does not utilize any cookies or collect any personal identifiable information. ImmunoFusion is free available in https://shiny.zhoulab.ac.cn/ImmunoFusion/.

### Dissecting the TME and quantifying cancer gene signatures

Consistent with our prior TME decomposition and cancer gene signature scoring methodology in TCCIA, we likewise employed IOBR^76^ which incorporates eight widely used open-source deconvolution methods. IOBR integrates a comprehensive compilation of 255 established cancer signatures. These signatures are categorized into three distinct groups: those associated with the TME, tumor metabolism, and tumor-intrinsic features. This organization enables a systematic investigation into the landscape of fusion genes and the immune microenvironment.

### Statistical analysis

We performed Kaplan-Meier survival analysis to generate and compare survival curves. The log-rank test was used for comparison. We also conducted multivariate survival analysis using the Cox regression model. All reported *P*-values are two-tailed, and a significance level of p≤0.05 was used unless otherwise specified. All statistical analyses and visualization were conducted using R v4.4.1.

## Acknowledgments

We are grateful for resources from the Bioinformatics Platform, Furong Laboratory and Bioinformatics Center, Xiangya Hospital, Central South University. This work was supported by the Guizhou Province College Students Innovation and Entrepreneurship Training Program (Grant No. ZHCX2023005), the Central South University Startup Funding, National Natural Science Foundation of China (Grant No. 82303953, 82060475), the Hunan Provincial Natural Science Foundation of China (Grant No. 2025JJ40079), Noncommunicable Chronic Diseases-National Science and Technology Major Project (Grant No. 2023ZD0502105), Chunhui program of the MOE (Ministry of Education in China) (Grant No. HZKY20220231), MOE Liberal arts and Social Sciences Foundation (Grant No. 24YJCZH462), Youth Science and Technology Elite Talent Project of Guizhou Provincial Department of Education (Grant No.QJJ-2024-333), Excellent Young Talent Cultivation Project of Zunyi City (Zunshi Kehe HZ (2023) 142), Future Science and Technology Elite Talent Cultivation Project of Zunyi Medical University (ZYSE 2023-02), and the Key Program of the Education Sciences Planning of Guizhou Province (Grant No.7).

## Contributions

YZ: Software, methodology, formal analysis, visualization, conceptualization, funding acquisition, writing original draft, review and editing. SW: Conceptualization, project administration, software, methodology, visualization, supervision, funding acquisition, writing original draft, review and editing. SL: Software, methodology, visualization. MC: Visualization. JGZ: Conceptualization, methodology, resources, supervision, funding acquisition, writing original draft, review and editing.

## Data availability

All relevant data reported in the study can be found in the article or on the ImmunoFusion website (accessible at https://shiny.zhoulab.ac.cn/ImmunoFusion/). For any other data requests, please contact the leader of this project Prof. Jian-Guo Zhou.

## Code availability

The fusion identification pipeline with an ensemble approach and corresponding scripts and logs for handling ImmunoFusion cohorts are available at https://github.com/OncoHarmony-Network/fusion-pipeline.

## Ethics statements

In this study, we conducted a secondary analysis of trial datasets, which was determined to carry minimal risk. The Institutional Review Board of the Second Affiliated Hospital, Zunyi Medical University (No. YXLL(KY-R)-2021-010) approved the study protocol. As per national legislation and institutional guidelines, written informed consent for participation was not deemed necessary for this particular study.

## References

1 Schram, A. M., Chang, M. T., Jonsson, P. & Drilon, A. Fusions in solid tumours: diagnostic strategies, targeted therapy, and acquired resistance. Nat Rev Clin Oncol 14, 735–748 (2017). 10.1038/nrclinonc.2017.127

2 Mitelman, F., Johansson, B. & Mertens, F. The impact of translocations and gene fusions on cancer causation. Nat Rev Cancer 7, 233–245 (2007).

3 Yoshihara, K. et al. The landscape and therapeutic relevance of cancer-associated transcript fusions. Oncogene 34, 4845–4854 (2015). 10.1038/onc.2014.406

4 Vogelstein, B. et al. Cancer genome landscapes. Science 339, 1546–1558 (2013). 10.1126/science.1235122

5 Babiceanu, M. et al. Recurrent chimeric fusion RNAs in non-cancer tissues and cells. Nucleic Acids Res 44, 2859–2872 (2016). 10.1093/nar/gkw032

6 Caldas, P. et al. Transcription readthrough is prevalent in healthy human tissues and associated with inherent genomic features. Commun Biol 7, 100 (2024). 10.1038/s42003-024-05779-5

7 Kim, D.-S. et al. CACG: a database for comparative analysis of conjoined genes. Genomics 100, 14–17 (2012). 10.1016/j.ygeno.2012.05.005

8 Ouedraogo, M. et al. The duplicated genes database: identification and functional annotation of co-localised duplicated genes across genomes. PLoS One 7, e50653 (2012). 10.1371/journal.pone.0050653

9 Lee, Y. A. et al. NTRK and RET fusion-directed therapy in pediatric thyroid cancer yields a tumor response and radioiodine uptake. J Clin Invest 131 (2021). 10.1172/JCI144847

10 Uhrig, S. et al. Accurate and efficient detection of gene fusions from RNA sequencing data. Genome Res 31, 448–460 (2021). 10.1101/gr.257246.119

11 Haas, B. J. et al. Accuracy assessment of fusion transcript detection via read-mapping and de novo fusion transcript assembly-based methods. Genome Biol 20, 213 (2019). 10.1186/s13059-019-1842-9

12 Hu, X. et al. TumorFusions: an integrative resource for cancer-associated transcript fusions. Nucleic Acids Res 46, D1144–D1149 (2018). 10.1093/nar/gkx1018

13 Jang, Y. E. et al. ChimerDB 4.0: an updated and expanded database of fusion genes. Nucleic Acids Res 48, D817–D824 (2020). 10.1093/nar/gkz1013

14 Gao, Q. et al. Driver Fusions and Their Implications in the Development and Treatment of Human Cancers. Cell Rep 23 (2018). 10.1016/j.celrep.2018.03.050

15 Davis, A. A. & Patel, V. G. The role of PD-L1 expression as a predictive biomarker: an analysis of all US Food and Drug Administration (FDA) approvals of immune checkpoint inhibitors. J Immunother Cancer 7, 278 (2019). 10.1186/s40425-019-0768-9

16 Yarchoan, M., Hopkins, A. & Jaffee, E. M. Tumor Mutational Burden and Response Rate to PD-1 Inhibition. N Engl J Med 377, 2500–2501 (2017). 10.1056/NEJMc1713444

17 Mazieres, J. et al. Immune checkpoint inhibitors for patients with advanced lung cancer and oncogenic driver alterations: results from the IMMUNOTARGET registry. Ann Oncol 30, 1321–1328 (2019). 10.1093/annonc/mdz167

18 Wu, Y.-M. et al. Inactivation of CDK12 Delineates a Distinct Immunogenic Class of Advanced Prostate Cancer. Cell 173 (2018). 10.1016/j.cell.2018.04.034

19 Creason, A. et al. A community challenge to evaluate RNA-seq, fusion detection, and isoform quantification methods for cancer discovery. Cell Syst 12 (2021). 10.1016/j.cels.2021.05.021

20 Apostolides, M. et al. MetaFusion: a high-confidence metacaller for filtering and prioritizing RNA-seq gene fusion candidates. Bioinformatics 37, 3144–3151 (2021). 10.1093/bioinformatics/btab249

21 Chen, Z. et al. TIGER: A Web Portal of Tumor Immunotherapy Gene Expression Resource. Genomics Proteomics Bioinformatics 21, 337–348 (2023). 10.1016/j.gpb.2022.08.004

22 Eddy, J. A. et al. CRI iAtlas: an interactive portal for immuno-oncology research. F1000Res 9, 1028 (2020). 10.12688/f1000research.25141.1

23 Huang, L. et al. FPIA: A database for gene fusion profiling and interactive analyses. Int J Cancer 150, 1504–1511 (2022). 10.1002/ijc.33921

24 Kim, P. et al. FusionGDB 2.0: fusion gene annotation updates aided by deep learning. Nucleic Acids Res 50, D1221–D1230 (2022). 10.1093/nar/gkab1056

25 Liu, Y. et al. IMPACT: A web server for exploring immunotherapeutic predictive and cancer prognostic biomarkers. Clin Transl Med 13, e1354 (2023). 10.1002/ctm2.1354

26 Sondka, Z. et al. COSMIC: a curated database of somatic variants and clinical data for cancer. Nucleic Acids Res 52, D1210–D1217 (2024). 10.1093/nar/gkad986

27 Wang, J. et al. A cloud-based resource for genome coordinate-based exploration and large-scale analysis of chromosome aberrations and gene fusions in cancer. Genes Chromosomes Cancer 62, 441–448 (2023). 10.1002/gcc.23128

28 Xia, Y. et al. ICBcomb: a comprehensive expression database for immune checkpoint blockade combination therapy. Brief Bioinform 25 (2023). 10.1093/bib/bbad457

29 Yang, J. et al. A pan-cancer immunogenomic atlas for immune checkpoint blockade immunotherapy. Cancer Res 82, 539–542 (2021). 10.1158/0008-5472.CAN-21-2335

30 Yang, M. et al. ICBatlas: A Comprehensive Resource for Depicting Immune Checkpoint Blockade Therapy Characteristics from Transcriptome Profiles. Cancer Immunol Res 10, 1398–1406 (2022). 10.1158/2326-6066.CIR-22-0249

31 Zhang, B. et al. Potential role of LPAR5 gene in prognosis and immunity of thyroid papillary carcinoma and pan-cancer. Sci Rep 13, 5850 (2023). 10.1038/s41598-023-32733-y

32 Greger, L. et al. Tandem RNA chimeras contribute to transcriptome diversity in human population and are associated with intronic genetic variants. PLoS One 9, e104567 (2014). 10.1371/journal.pone.0104567

33 Tate, J. G. et al. COSMIC: the Catalogue Of Somatic Mutations In Cancer. Nucleic Acids Res 47, D941–D947 (2019). 10.1093/nar/gky1015

34 Arafeh, R., Shibue, T., Dempster, J. M., Hahn, W. C. & Vazquez, F. The present and future of the Cancer Dependency Map. Nat Rev Cancer 25, 59–73 (2025). 10.1038/s41568-024-00763-x

35 Klijn, C. et al. A comprehensive transcriptional portrait of human cancer cell lines. Nat Biotechnol 33, 306–312 (2015). 10.1038/nbt.3080

36 Lee, M. et al. ChimerDB 3.0: an enhanced database for fusion genes from cancer transcriptome and literature data mining. Nucleic Acids Res 45, D784–D789 (2017). 10.1093/nar/gkw1083

37 Dehghannasiri, R. et al. Improved detection of gene fusions by applying statistical methods reveals oncogenic RNA cancer drivers. Proc Natl Acad Sci U S A 116, 15524–15533 (2019). 10.1073/pnas.1900391116

38 Alaei-Mahabadi, B., Bhadury, J., Karlsson, J. W., Nilsson, J. A. & Larsson, E. Global analysis of somatic structural genomic alterations and their impact on gene expression in diverse human cancers. Proc Natl Acad Sci U S A 113, 13768–13773 (2016).

39 Integrated genomic analyses of ovarian carcinoma. Nature 474, 609–615 (2011). 10.1038/nature10166

40 Comprehensive molecular characterization of gastric adenocarcinoma. Nature 513, 202–209 (2014). 10.1038/nature13480

41 Hafstað, V. et al. Improved detection of clinically relevant fusion transcripts in cancer by machine learning classification. BMC Genomics 24, 783 (2023). 10.1186/s12864-023-09889-y

42 Sung, C. et al. Integrative analysis of risk factors for immune-related adverse events of checkpoint blockade therapy in cancer. Nat Cancer 4, 844–859 (2023). 10.1038/s43018-023-00572-5

43 Beroukhim, R. et al. The landscape of somatic copy-number alteration across human cancers. Nature 463, 899–905 (2010). 10.1038/nature08822

44 Su, C. Y. et al. MTAP is an independent prognosis marker and the concordant loss of MTAP and p16 expression predicts short survival in non-small cell lung cancer patients. Eur J Surg Oncol 40, 1143–1150 (2014). 10.1016/j.ejso.2014.04.017

45 Han, G. et al. 9p21 loss confers a cold tumor immune microenvironment and primary resistance to immune checkpoint therapy. Nat Commun 12, 5606 (2021). 10.1038/s41467-021-25894-9

46 Keyel, P. A. et al. Methylthioadenosine reprograms macrophage activation through adenosine receptor stimulation. PLoS One 9, e104210 (2014). 10.1371/journal.pone.0104210

47 Limm, K., Wallner, S., Milenkovic, V. M., Wetzel, C. H. & Bosserhoff, A.-K. The metabolite 5’-methylthioadenosine signals through the adenosine receptor A2B in melanoma. Eur J Cancer 50, 2714–2724 (2014). 10.1016/j.ejca.2014.07.005

48 Munshi, R., Clanachan, A. S. & Baer, H. P. 5’-Deoxy-5’-methylthioadenosine: a nucleoside which differentiates between adenosine receptor types. Biochem Pharmacol 37, 2085–2089 (1988).

49 Barekatain, Y. et al. Homozygous MTAP deletion in primary human glioblastoma is not associated with elevation of methylthioadenosine. Nat Commun 12, 4228 (2021). 10.1038/s41467-021-24240-3

50 Wells, A. D. & Morawski, P. A. New roles for cyclin-dependent kinases in T cell biology: linking cell division and differentiation. Nat Rev Immunol 14, 261–270 (2014). 10.1038/nri3625

51 Patil, N. S. et al. Intratumoral plasma cells predict outcomes to PD-L1 blockade in non-small cell lung cancer. Cancer Cell 40 (2022). 10.1016/j.ccell.2022.02.002

52 Tomlins, S. A. et al. Distinct classes of chromosomal rearrangements create oncogenic ETS gene fusions in prostate cancer. Nature 448, 595–599 (2007).

53 Sanderson, S. M., Mikhael, P. G., Ramesh, V., Dai, Z. & Locasale, J. W. Nutrient availability shapes methionine metabolism in p16/MTAP-deleted cells. Sci Adv 5, eaav7769 (2019). 10.1126/sciadv.aav7769

54 Carroll, T. M. et al. Tumor monocyte content predicts immunochemotherapy outcomes in esophageal adenocarcinoma. Cancer Cell 41 (2023). 10.1016/j.ccell.2023.06.006

55 Creelan, B. C. et al. Tumor-infiltrating lymphocyte treatment for anti-PD-1-resistant metastatic lung cancer: a phase 1 trial. Nat Med 27, 1410–1418 (2021). 10.1038/s41591-021-01462-y

56 Wang, S. et al. TCCIA: a comprehensive resource for exploring CircRNA in cancer immunotherapy. J Immunother Cancer 12 (2024). 10.1136/jitc-2023-008040

57 Gide, T. N. et al. Distinct Immune Cell Populations Define Response to Anti-PD-1 Monotherapy and Anti-PD-1/Anti-CTLA-4 Combined Therapy. Cancer Cell 35 (2019). 10.1016/j.ccell.2019.01.003

58 Grasso, C. S. et al. Conserved Interferon-γ Signaling Drives Clinical Response to Immune Checkpoint Blockade Therapy in Melanoma. Cancer Cell 38 (2020). 10.1016/j.ccell.2020.08.005

59 Motzer, R. J. et al. Molecular Subsets in Renal Cancer Determine Outcome to Checkpoint and Angiogenesis Blockade. Cancer Cell 38 (2020). 10.1016/j.ccell.2020.10.011

60 Campbell, K. M. et al. Prior anti-CTLA-4 therapy impacts molecular characteristics associated with anti-PD-1 response in advanced melanoma. Cancer Cell 41 (2023). 10.1016/j.ccell.2023.03.010

61 Hugo, W. et al. Genomic and Transcriptomic Features of Response to Anti-PD-1 Therapy in Metastatic Melanoma. Cell 165, 35–44 (2016). 10.1016/j.cell.2016.02.065

62 Riaz, N. et al. Tumor and Microenvironment Evolution during Immunotherapy with Nivolumab. Cell 171 (2017). 10.1016/j.cell.2017.09.028

63 Liu, S. et al. Response and recurrence correlates in individuals treated with neoadjuvant anti-PD-1 therapy for resectable oral cavity squamous cell carcinoma. Cell Rep Med 2, 100411 (2021). 10.1016/j.xcrm.2021.100411

64 Bellmunt, J. et al. Adjuvant atezolizumab versus observation in muscle-invasive urothelial carcinoma (IMvigor010): a multicentre, open-label, randomised, phase 3 trial. Lancet Oncol 22, 525–537 (2021). 10.1016/S1470-2045(21)00004-8

65 Berger, M. F. et al. Melanoma genome sequencing reveals frequent PREX2 mutations. Nature 485, 502–506 (2012). 10.1038/nature11071

66 Tumeh, P. C. et al. PD-1 blockade induces responses by inhibiting adaptive immune resistance. Nature 515, 568–571 (2014). 10.1038/nature13954

67 Mariathasan, S. et al. TGFβ attenuates tumour response to PD-L1 blockade by contributing to exclusion of T cells. Nature 554, 544–548 (2018). 10.1038/nature25501

68 Roper, N. et al. Notch signaling and efficacy of PD-1/PD-L1 blockade in relapsed small cell lung cancer. Nat Commun 12, 3880 (2021). 10.1038/s41467-021-24164-y

69 Cui, C. et al. Ratio of the interferon-γ signature to the immunosuppression signature predicts anti-PD-1 therapy response in melanoma. NPJ Genom Med 6, 7 (2021). 10.1038/s41525-021-00169-w

70 McDermott, D. F. et al. Clinical activity and molecular correlates of response to atezolizumab alone or in combination with bevacizumab versus sunitinib in renal cell carcinoma. Nat Med 24, 749–757 (2018). 10.1038/s41591-018-0053-3

71 Kim, S. T. et al. Comprehensive molecular characterization of clinical responses to PD-1 inhibition in metastatic gastric cancer. Nat Med 24, 1449–1458 (2018). 10.1038/s41591-018-0101-z

72 Auslander, N., et al. Publisher Correction: Robust prediction of response to immune checkpoint blockade therapy in metastatic melanoma. Nat Med 24, 1942 (2018). 10.1038/s41591-018-0247-8

73 Cloughesy, T. F. et al. Neoadjuvant anti-PD-1 immunotherapy promotes a survival benefit with intratumoral and systemic immune responses in recurrent glioblastoma. Nat Med 25, 477–486 (2019). 10.1038/s41591-018-0337-7

74 (!!! INVALID CITATION !!!).

75 Li, S. et al. Facilitating integrative and personalized oncology omics analysis with UCSCXenaShiny. Commun Biol 7, 1200 (2024). 10.1038/s42003-024-06891-2

76 Zeng, D. et al. IOBR: Multi-Omics Immuno-Oncology Biological Research to Decode Tumor Microenvironment and Signatures. Front Immunol 12, 687975 (2021). 10.3389/fimmu.2021.687975

