## Supplementary_Figures for "ImmunoFusion: A Unified Platform for Investigating RNA-seq-Derived Gene Fusions in Cancer and Immunotherapy"

### Supplementary Figure 1

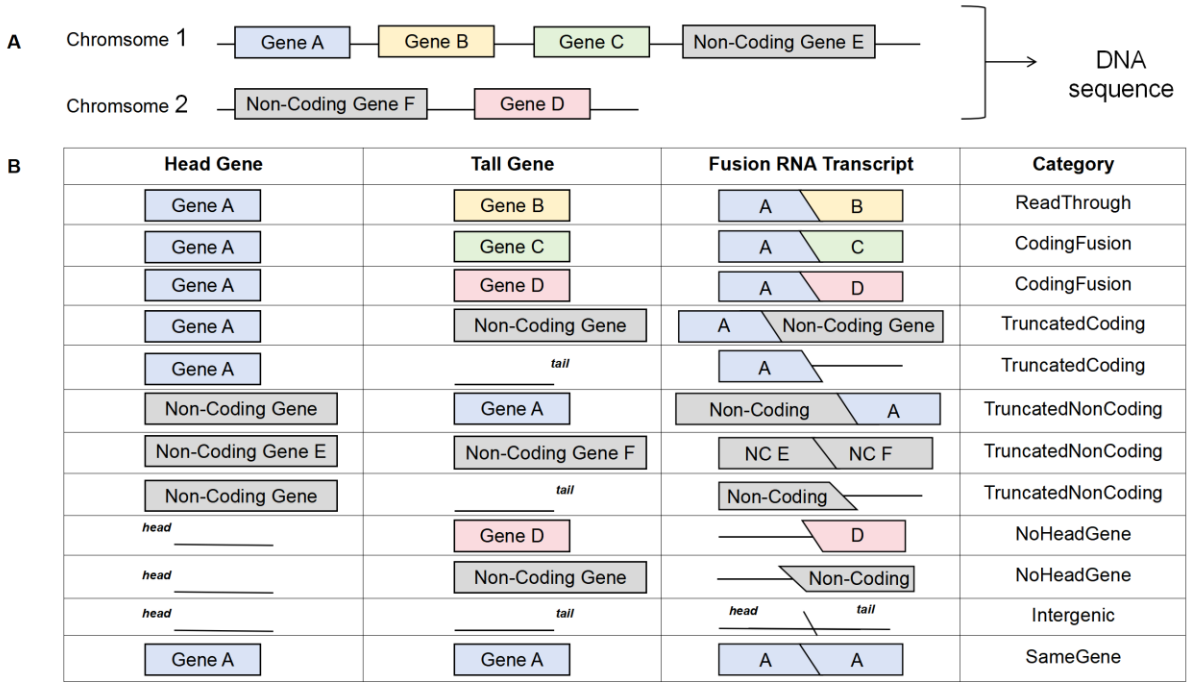


**Supplementary Figure 1. MetaFusion-defined RNA-seq gene fusion categories.** (**A**) Two chromosomes with coding (colored) and non-coding (grey) genes, intergenic sequences shown as black lines. (**B**) Head and tail genes from DNA in (A), resulting RNA fusion transcripts, and their assigned fusion categories. Abbreviation: NC, non-coding.

#### Supplementary Figure 2

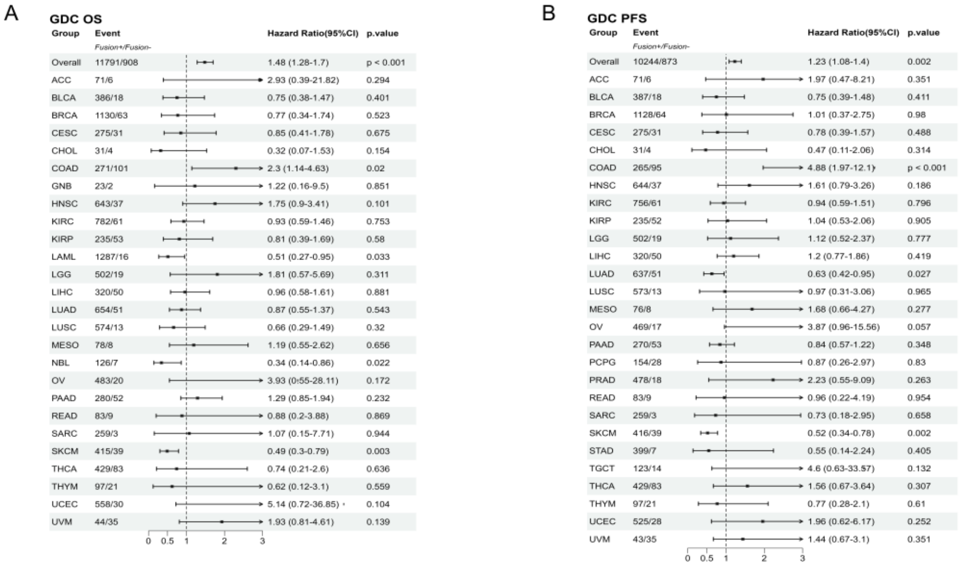


**Supplementary Figure 2. Prognostic impact of gene fusions in pan-cancer GDC cohorts.** (**A,B**) Forest plots showing hazard ratios (HR) and 95% confidence intervals (CI) for (**A**) overall survival (OS) and (**B**) progression-free survival (PFS) from tumor-type-stratified Cox proportional hazards models. Analysis includes GDC cohorts (including TCGA, TARGET, and CPTAC; OS N=12,699; PFS N=11,117) without adjustments. “Overall” represents the aggregated results of all cohorts. p<0.001 are marked as ‘p<0.001’ instead of a detailed value.

#### Supplementary Figure 3

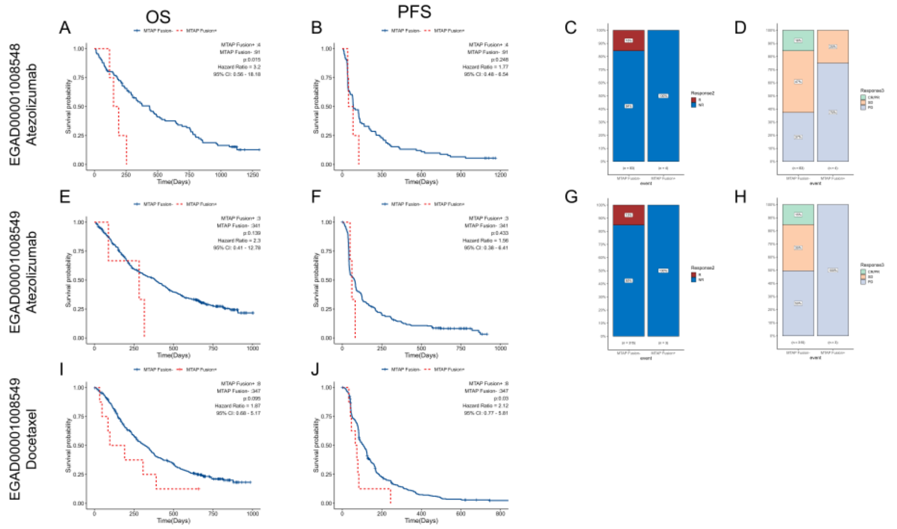


**Supplementary Figure 3. Prognostic impact of MTAP fusion status in NSCLC patients treated with atezolizumab or docetaxel.** (**A,B**) Kaplan-Meier curves for (**A**) overall survival (OS) and (**B**) progression-free survival (PFS) in the EGAD00001008548 (POPLAR) anti-PD-L1 (atezolizumab) subgroup (N for total=84, N for Fusion+: 4), stratified by MTAP fusion status (Fusion+: red; Fusion-: blue). (**E,F**) EGAD00001008549 (OAK) anti-PD-L1 (atezolizumab) subgroup (N=318, Fusion+: 3). **(I,J)** OAK chemotherapy (docetaxel) subgroup (N=347, Fusion+: 8). The best overall responses (BOR) (RECIST 1.1) were compared between patient groups stratified by MTAP fusion detected. Response (R; red): CR/PR (green), complete or partial response; nonresponse (NR; blue): SD (orange), stable disease; PD (blue) progressive disease. *P*-values and odds ratios (OR) were calculated using two-sided Fisher's Exact tests. (**C,G**) Response rates (R vs NR) were compared between the two groups. (**C**) POPLAR p = 1. (**G**) OAK p = 1. (**D,H**) Percentages of PD were compared between the two groups. (**D**) POPLAR p = 0.29, OR = 4.94. (**H**) OAK p = 0.25. Note: POPLAR docetaxel subgroup excluded due to insufficient MTAP fusion-positive cases (n=1).
