## Supplementary_Methods for "ImmunoFusion: A Unified Platform for Investigating RNA-seq-Derived Gene Fusions in Cancer and Immunotherapy"

**Systematic search of bulk RNA-seq data from ICB cohorts**

Referring to our previous cohort collection process in TCCIA^1^, we performed a systematic search using PubMed (<https://www.ncbi.nlm.nih.gov/pubmed/>) to identify articles related to bulk RNA-seq data from solid cancer patients treated with immune-checkpoint blockade (ICB). In total, we collected 29 studies^2-28^ related to ICB that provided raw RNA-seq datasets. We gathered relevant clinical data from publications and clinical meta documents associated with these RNA-seq datasets. Additionally, we extracted and validated information on the cohorts’ fundamental characteristics, such as sample size, treatment methods, and specific drugs used, based on the abstracts and supplementary files from original manuscripts. It should be noted that, for Patil *et al*.^16^ study, two clinical cohorts were included; and for the study identified as PHS000452, the two patient subgroups had distinct drug treatments and clinical annotations. Hence, we treated them as two separate cohorts during the analysis, and for Sung *et al.*^14^, we considered this cohort as pan-cancer rather than specific to any single cancer type, because it compiled germline exomes and blood transcriptomes along with clinical data. In summary, RNA-seq derived gene fusions, treatment response, and clinical benefits were analyzed for twelve cancer types in our study. Our analysis included over 4,647 clinical samples from 29 cohorts treated with ICBs such as PD-1/PD-L1 and CTLA-4 inhibitors, as well as other treatments. The analysis considered both pre-treatment and on-treatment responses.

**Gene fusion integration and evaluation**

We used a modified version of MetaFusion^29^ for integration of identified gene fusion candidates from Arriba^30^ and Star-Fusion^31^. This process includes: 1) converts fusion calls from different identification methods to Common Fusion Format (CFF), a novel file type that it has developed to standardize fusion caller outputs; 2) employs both breakpoint and gene name intersections to determine whether multiple calls should be consolidated into a single event; 3) adopts the filtering approach referenced from Arriba to generate a blocklist, which we have incorporated as the blocklist column in our integration results (this enables researchers to filter out fusions by their own purposes); 4) categorizes fusions into one of seven classes based on the coding status of the fusion partners and their genomic proximity (Supplementary Figure 1). Subsequently, FusionAnnotator (v0.2.0)^31^ was used to generate the cancer_db_hits column for annotating fusions, flagging those previously reported as relevant to cancer biology, as well as identifying paralogs, promiscuity, and potential red herrings (fusion pairs that may not be relevant to cancer and are potential false positives). As the official MetaFusion Docker image uses GRCh37 GTF reference files, we generated GRCh38 GTF reference files using their provided method and created a corresponding Docker image (<https://github.com/users/ShixiangWang/packages/container/package/metafusion2>).

During our manual inspection of the integrated fusion results, we identified numerous fusion pairs in TCGA tumor sample data that were not reported in previous studies^31-36^. This is primarily because prior research tends to focus on reporting highly confident positive results. Additionally, these unreported fusions could result from alignment artifacts^30^, low-quality RNA^37^, or transcript variants also present in healthy tissues^38^. Therefore, drawing on prior studies^30-32,39-41^, we have proposed a two-step scoring strategy to evaluate the credibility of fusions.

- For normal samples, fusion events are scored as follows: 1) If a fusion event is listed in the Arriba blocklist, it receives a score of 0, which remains unchanged in the following steps; 2) If a fusion event has max_split_cnt ≥ 1 and max_span_cnt ≥ 1, it gains +1 to its score; 3) If both Arriba and STAR-Fusion detect the same fusion event, an additional +1 is added; 4) If FusionAnnotator previously annotated the event as a normal fusion, +2 is added to the score. A fusion from normal samples with score ≥ 2 is considered as confident.
- For tumor samples, fusion events are scored as follows: 1) If a fusion event is listed in the Arriba blocklist, it receives a score of 0, which remains unchanged in the following steps; 2) If a fusion event is detected as a confident normal fusion (i.e., score ≥ 2) in our analysis, it is assigned a score of 0, which remains unchanged; 3) If a fusion event has max_split_cnt ≥ 1 and max_span_cnt ≥ 1, while with at least one value < 5, it gains +1 to its score; if both values are ≥ 5, it gains +2; 4) If both Arriba and STAR-Fusion detect the same fusion event, an additional +1 is added. 5) If FusionAnnotator previously annotated the event as a tumor fusion, +2 is added to the score. Similarly, A fusion from tumor samples with score ≥ 2 is considered as confident.

**Clinical and IOBR/TME data preprocessing**

We obtained clinical data from the Genomic Data Commons (GDC) via TCGAbiolinks^42^ and UCSC Xena^43^ via UCSCXenaShiny^44^. Subsequently, we performed data cleaning and renaming operations, which included filling missing values, removing duplicates, renaming columns, and processing the gender field. We merged survival data, purity data, genomic instability data, and subtype data based on the Sample ID. We verified that the majority of valid rows in the clinical data provided by GDC matched those in the clinical data provided by UCSCXenaShiny. Subsequently, we used the clinical information organized by UCSCXenaShiny to fill in the missing values in the GDC data, merging them based on the Sample ID field.

For sample filtering: 1) We excluded the clinical information of samples for which GDC did not provide fastq files. 2) For regarding composite samples, In the TARGET database, samples such as TARGET-30-PASYPX-01A and TARGET-30-PAIXIF-01A, which are mixtures of samples from different patients, were removed from our dataset due to their limited quantity and the presence of individual TARGET-30-PASYPX-01A samples. In the CPTAC database, composite samples like C3L-03400-05 and C3L-03400-02, which consist of multiple samples from the same patient and are more numerous, were retained. In the TCGA database did not exhibit this issue; 3) For cohorts missing clinical information for some samples (PRJEB25780, PRJNA356761 and PRJNA498500, including samples like “SRR5088919” or others), we removed the corresponding samples in required analysis.

For cohort naming: We referred to the TCGA tumor type naming conventions and created the ImmunoFusion cohort naming system based on tumor type information and data sources. We divided the three major databases into 50 cohorts. For specific cases, such as AML in TARGET and HGSC in CPTAC, we named them TARGET-LAML and CPTAC-OV, respectively, since TCGA typically names these tumor types as LAML and OV. Samples derived from normal samples were categorized into CPTAC-NORMAL, TARGET-NORMAL, and TCGA-NORMAL cohorts. Tumor cell line samples from the TARGET database were placed in a separate TARGET-CELL cohort. The complete mapping table of public database naming and source information is available at Supplementary Table3 and ([https://oncoharmony-network.github.io/doc-immunofusion/methods.html#immunofusion-cohort-naming-maping](https://oncoharmony-network.github.io/doc-immunofusion/methods.html" \l "immunofusion-cohort-naming-maping)). ICB cohorts, such as NSCLC and SCLC, retained their original cancer types, as they lacked specific cancer type classifications like LUSC.

For IOBR/TME data: Following the GDC barcode naming rules (<https://docs.gdc.cancer.gov/Encyclopedia/pages/TCGA_Barcode/>), we selected rows with later Portion numbers for the same sample to obtain more accurate immune microenvironment information. For matching purposes, we used the portion of the barcode before Portion as the sample name. Ultimately, we generated three well-processed data frames. The complete processing code is available at (<https://oncoharmony-network.github.io/doc-immunofusion/methods.html>).

**References**

1	Wang, S. *et al.* TCCIA: a comprehensive resource for exploring CircRNA in cancer immunotherapy. *J Immunother Cancer* **12** (2024). <https://doi.org/10.1136/jitc-2023-008040>

2	Bellmunt, J. *et al.* Adjuvant atezolizumab versus observation in muscle-invasive urothelial carcinoma (IMvigor010): a multicentre, open-label, randomised, phase 3 trial. *Lancet Oncol* **22**, 525-537 (2021). <https://doi.org/10.1016/S1470-2045(21)00004-8>

3	McDermott, D. F. *et al.* Clinical activity and molecular correlates of response to atezolizumab alone or in combination with bevacizumab versus sunitinib in renal cell carcinoma. *Nat Med* **24**, 749-757 (2018). <https://doi.org/10.1038/s41591-018-0053-3>

4	Kim, S. T. *et al.* Comprehensive molecular characterization of clinical responses to PD-1 inhibition in metastatic gastric cancer. *Nat Med* **24**, 1449-1458 (2018). <https://doi.org/10.1038/s41591-018-0101-z>

5	Grasso, C. S. *et al.* Conserved Interferon-γ Signaling Drives Clinical Response to Immune Checkpoint Blockade Therapy in Melanoma. *Cancer Cell* **38** (2020). <https://doi.org/10.1016/j.ccell.2020.08.005>

6	Gide, T. N. *et al.* Distinct Immune Cell Populations Define Response to Anti-PD-1 Monotherapy and Anti-PD-1/Anti-CTLA-4 Combined Therapy. *Cancer Cell* **35** (2019). <https://doi.org/10.1016/j.ccell.2019.01.003>

7	Liu, D. *et al.* Evolution of delayed resistance to immunotherapy in a melanoma responder. *Nat Med* **27**, 985-992 (2021). <https://doi.org/10.1038/s41591-021-01331-8>

8	Snyder, A. *et al.* Genetic basis for clinical response to CTLA-4 blockade in melanoma. *N Engl J Med* **371**, 2189-2199 (2014). <https://doi.org/10.1056/NEJMoa1406498>

9	Hugo, W. *et al.* Genomic and Transcriptomic Features of Response to Anti-PD-1 Therapy in Metastatic Melanoma. *Cell* **165**, 35-44 (2016). <https://doi.org/10.1016/j.cell.2016.02.065>

10	Miao, D. *et al.* Genomic correlates of response to immune checkpoint therapies in clear cell renal cell carcinoma. *Science* **359**, 801-806 (2018). <https://doi.org/10.1126/science.aan5951>

11	Zhao, J. *et al.* Immune and genomic correlates of response to anti-PD-1 immunotherapy in glioblastoma. *Nat Med* **25**, 462-469 (2019). <https://doi.org/10.1038/s41591-019-0349-y>

12	Rosenbaum, E. *et al.* Immune-related Adverse Events after Immune Checkpoint Blockade-based Therapy Are Associated with Improved Survival in Advanced Sarcomas. *Cancer Res Commun* **3**, 2118-2125 (2023). <https://doi.org/10.1158/2767-9764.CRC-22-0140>

13	Gettinger, S. *et al.* Impaired HLA Class I Antigen Processing and Presentation as a Mechanism of Acquired Resistance to Immune Checkpoint Inhibitors in Lung Cancer. *Cancer Discov* **7**, 1420-1435 (2017). <https://doi.org/10.1158/2159-8290.CD-17-0593>

14	Sung, C. *et al.* Integrative analysis of risk factors for immune-related adverse events of checkpoint blockade therapy in cancer. *Nat Cancer* **4**, 844-859 (2023). <https://doi.org/10.1038/s43018-023-00572-5>

15	Patil, N. S. *et al.* Intratumoral plasma cells predict outcomes to PD-L1 blockade in non-small cell lung cancer. *Cancer Cell* **40** (2022). <https://doi.org/10.1016/j.ccell.2022.02.002>

16	Berger, M. F. *et al.* Melanoma genome sequencing reveals frequent PREX2 mutations. *Nature* **485**, 502-506 (2012). <https://doi.org/10.1038/nature11071>

17	Motzer, R. J. *et al.* Molecular Subsets in Renal Cancer Determine Outcome to Checkpoint and Angiogenesis Blockade. *Cancer Cell* **38** (2020). <https://doi.org/10.1016/j.ccell.2020.10.011>

18	Cloughesy, T. F. *et al.* Neoadjuvant anti-PD-1 immunotherapy promotes a survival benefit with intratumoral and systemic immune responses in recurrent glioblastoma. *Nat Med* **25**, 477-486 (2019). <https://doi.org/10.1038/s41591-018-0337-7>

19	Vos, J. L. *et al.* Nivolumab plus ipilimumab in advanced salivary gland cancer: a phase 2 trial. *Nat Med* **29**, 3077-3089 (2023). <https://doi.org/10.1038/s41591-023-02518-x>

20	Roper, N. *et al.* Notch signaling and efficacy of PD-1/PD-L1 blockade in relapsed small cell lung cancer. *Nat Commun* **12**, 3880 (2021). <https://doi.org/10.1038/s41467-021-24164-y>

21	Tumeh, P. C. *et al.* PD-1 blockade induces responses by inhibiting adaptive immune resistance. *Nature* **515**, 568-571 (2014). <https://doi.org/10.1038/nature13954>

22	Nguyen, V. P. *et al.* A Pilot Study of Neoadjuvant Nivolumab, Ipilimumab, and Intralesional Oncolytic Virotherapy for HER2-negative Breast Cancer. *Cancer Res Commun* **3**, 1628-1637 (2023). <https://doi.org/10.1158/2767-9764.CRC-23-0145>

23	Campbell, K. M. *et al.* Prior anti-CTLA-4 therapy impacts molecular characteristics associated with anti-PD-1 response in advanced melanoma. *Cancer Cell* **41** (2023). <https://doi.org/10.1016/j.ccell.2023.03.010>

24	Auslander, N. *et al.* Publisher Correction: Robust prediction of response to immune checkpoint blockade therapy in metastatic melanoma. *Nat Med* **24**, 1942 (2018). <https://doi.org/10.1038/s41591-018-0247-8>

25	Cui, C. *et al.* Ratio of the interferon-γ signature to the immunosuppression signature predicts anti-PD-1 therapy response in melanoma. *NPJ Genom Med* **6**, 7 (2021). <https://doi.org/10.1038/s41525-021-00169-w>

26	Liu, S. *et al.* Response and recurrence correlates in individuals treated with neoadjuvant anti-PD-1 therapy for resectable oral cavity squamous cell carcinoma. *Cell Rep Med* **2**, 100411 (2021). <https://doi.org/10.1016/j.xcrm.2021.100411>

27	Mariathasan, S. *et al.* TGFβ attenuates tumour response to PD-L1 blockade by contributing to exclusion of T cells. *Nature* **554**, 544-548 (2018). <https://doi.org/10.1038/nature25501>

28	Riaz, N. *et al.* Tumor and Microenvironment Evolution during Immunotherapy with Nivolumab. *Cell* **171** (2017). <https://doi.org/10.1016/j.cell.2017.09.028>

29	Apostolides, M. *et al.* MetaFusion: a high-confidence metacaller for filtering and prioritizing RNA-seq gene fusion candidates. *Bioinformatics* **37**, 3144-3151 (2021). <https://doi.org/10.1093/bioinformatics/btab249>

30	Uhrig, S. *et al.* Accurate and efficient detection of gene fusions from RNA sequencing data. *Genome Res* **31**, 448-460 (2021). <https://doi.org/10.1101/gr.257246.119>

31	Haas, B. J. *et al.* Accuracy assessment of fusion transcript detection via read-mapping and de novo fusion transcript assembly-based methods. *Genome Biol* **20**, 213 (2019). <https://doi.org/10.1186/s13059-019-1842-9>

32	Gao, Q. *et al.* Driver Fusions and Their Implications in the Development and Treatment of Human Cancers. *Cell Rep* **23** (2018). <https://doi.org/10.1016/j.celrep.2018.03.050>

33	Yoshihara, K. *et al.* The landscape and therapeutic relevance of cancer-associated transcript fusions. *Oncogene* **34**, 4845-4854 (2015). <https://doi.org/10.1038/onc.2014.406>

34	Hu, X. *et al.* TumorFusions: an integrative resource for cancer-associated transcript fusions. *Nucleic Acids Res* **46**, D1144-D1149 (2018). <https://doi.org/10.1093/nar/gkx1018>

35	Dehghannasiri, R. *et al.* Improved detection of gene fusions by applying statistical methods reveals oncogenic RNA cancer drivers. *Proc Natl Acad Sci U S A* **116**, 15524-15533 (2019). <https://doi.org/10.1073/pnas.1900391116>

36	Alaei-Mahabadi, B., Bhadury, J., Karlsson, J. W., Nilsson, J. A. & Larsson, E. Global analysis of somatic structural genomic alterations and their impact on gene expression in diverse human cancers. *Proc Natl Acad Sci U S A* **113**, 13768-13773 (2016).

37	Cheng, W. *et al.* CeO2/MXene heterojunction-based ultrasensitive electrochemiluminescence biosensing for BCR-ABL fusion gene detection combined with dual-toehold strand displacement reaction for signal amplification. *Biosens Bioelectron* **210**, 114287 (2022). <https://doi.org/10.1016/j.bios.2022.114287>

38	Babiceanu, M. *et al.* Recurrent chimeric fusion RNAs in non-cancer tissues and cells. *Nucleic Acids Res* **44**, 2859-2872 (2016). <https://doi.org/10.1093/nar/gkw032>

39	Shah, J. B. *et al.* Analysis of matched primary and recurrent BRCA1/2 mutation-associated tumors identifies recurrence-specific drivers. *Nat Commun* **13**, 6728 (2022). <https://doi.org/10.1038/s41467-022-34523-y>

40	Hayette, S. *et al.* Performances of Targeted RNA Sequencing for the Analysis of Fusion Transcripts, Gene Mutation, and Expression in Hematological Malignancies. *Hemasphere* **5**, e522 (2021). <https://doi.org/10.1097/HS9.0000000000000522>

41	Mariella, E. *et al.* Transcriptome-wide gene expression outlier analysis pinpoints therapeutic vulnerabilities in colorectal cancer. *Mol Oncol* **18**, 1460-1485 (2024). <https://doi.org/10.1002/1878-0261.13622>

42	Colaprico, A. *et al.* TCGAbiolinks: an R/Bioconductor package for integrative analysis of TCGA data. *Nucleic Acids Res* **44**, e71 (2016). <https://doi.org/10.1093/nar/gkv1507>

43	Goldman, M. J. *et al.* Visualizing and interpreting cancer genomics data via the Xena platform. *Nat Biotechnol* **38**, 675-678 (2020). <https://doi.org/10.1038/s41587-020-0546-8>

44	Li, S. *et al.* Facilitating integrative and personalized oncology omics analysis with UCSCXenaShiny. *Commun Biol* **7**, 1200 (2024). <https://doi.org/10.1038/s42003-024-06891-2>
